# A cold-adapted Cas9 nuclease from the deep ocean

**DOI:** 10.64898/2026.10.14.759226

**Authors:** Alba Olaso-Llorca, Francesco Colizzi, Gerard Mingarro, Pablo Sánchez, Carlos Sánchez-Bolaños, Constantino Andrés Molina, Andrea Martínez-Cazorla, Rafael Serrano-Durán, Tomas Urbaitis, Natalia Rodrigo Melero, Albert Bosch, María Magallon-Mosella, Mark Margitfalvi, Migle Stitilyte, Carlo Carolis, Damià Romero-Moya, Alessandra Giorgetti, Andrés Velázquez-Mudarra, Dejan Rajkovskii, F. Del Caño-Ochoa, Diana López-Farfán, Javier Gómez, Alfonso Ferrández-Roldán, Santiago Ramón-Maiques, Elena Gómez Díaz, Cristian Cañestro, Carlos M. Duarte, Josep M. Gasol, Francisco J.M. Mojica, Antonio Sánchez-Amat, Miguel Ángel Moreno-Mateos, Giedrius Gasiunas, Silvia G. Acinas, Julián Cerón

## Abstract

The abundance of CRISPR systems in bacteria and archaea fuels the search for Cas nucleases that perform where established enzymes fail. Deep Ocean 1 Cas9 (DO1Cas9) is to our knowledge the first Cas9 nuclease characterized from the deep ocean, recovered from a metagenomic sample at 1,500 m depth in the Atlantic Ocean. DO1Cas9 shows cold-adapted structural flexibility, thermal susceptibility, salt-dependent stability, higher specificity than SpCas9 and cleaves double-stranded DNA efficiently under cold conditions where mesophilic SpCas9 retains only residual activity. DO1Cas9 edits genomes in diverse organisms such as *Caenorhabditis elegans*, zebrafish, the marine tunicate *Oikopleura dioica*, and the malaria parasite *Plasmodium falciparum*. In the marine bacterium *Marinomonas mediterranea*, DO1Cas9 programmed against an invading phage reduced plaquing efficiency about 10-fold more than SpCas9. These results expand the CRISPR-Cas9 toolbox with a nuclease for cold-chain editing and marine biotechnology.

---

Over the last decade, nucleases within the CRISPR-Cas toolbox have expanded mainly by engineering variants of existing Cas nucleases or by mining environmental samples or microbiomes for new Cas proteins ^1,2^. The estimation of the existence of millions of bacterial and archaeal species^3^ implies a vast number of CRISPR-Cas systems mediating adaptive immunity that remain to be uncovered and described, particularly in environments of difficult access, such as the deep ocean. The analyses of global ocean metagenomic datasets, including samples of vertical profiles, generated during the scientific circumnavigation of the Malaspina expedition^4,5^, identified 48 novel Cas9 proteins^6^. Among them, Deep Ocean 1 Cas9 (DO1Cas9) was identified in a sample collected at 1500 meters of depth in the Atlantic Ocean with an average temperature about 2-4°C^7^. DO1Cas9 is a Cas9 of 991 amino acids. Single-guide RNAs (sgRNAs) were designed from the DO1Cas9-specific crRNA (36 nt) and tracrRNA (111 nt) identified at its CRISPR-Cas locus (**Fig. 1a and Supplementary Data 1**). In vitro-generated DO1Cas9 and its sgRNA were used to interrogate a target plasmid library containing a randomized 7N PAM (Protospacer Adjacent Motif) sequence^8^ (**Fig. 1b**). Sequencing of cleaved plasmids showed that the predominant cut site was located 3 bp upstream of the PAM, generating blunt ends, and defined the PAM as 5^′^-NRVDHCN-3^′^. Further analysis by in vitro digestion with purified DO1Cas9 protein expanded the potential PAM permissiveness to 5^′^-NVNNNCN-3^′^ (**Fig. 1c and Supplementary Data 2**).

**Figure 1.**
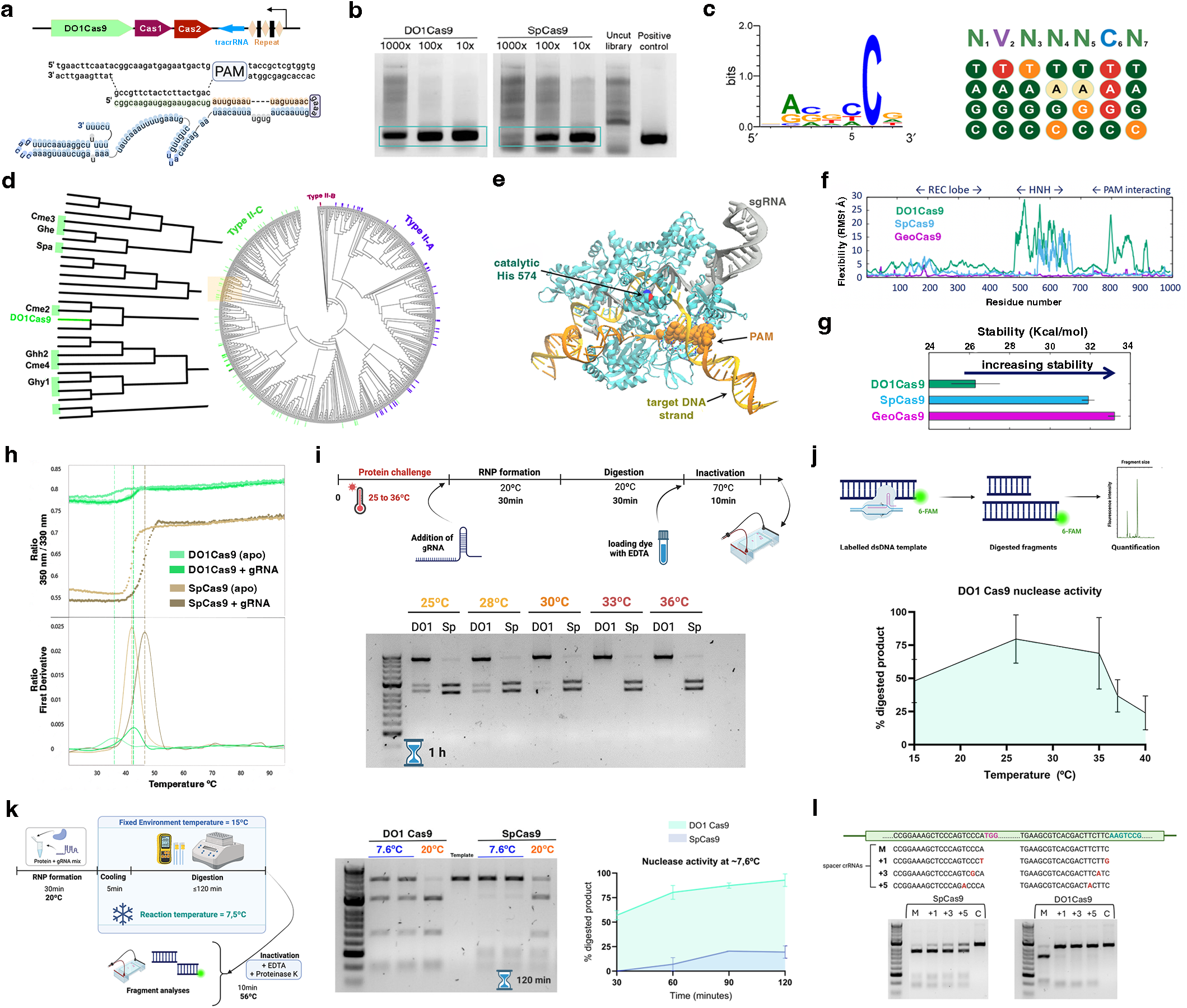
Description and cold-adapted properties of DO1Cas9. **a**, Top: DO1Cas9 CRISPR–Cas locus. Bottom: RNA secondary structure (predicted using trRosettaRNA^24^) and sequence of a DO1Cas9 sgRNA, consisting of a 20-nt variable region (green) matching a target sequence upstream of the PAM, followed by repeat (orange) and tracrRNA (blue) sequences connected by a linker (purple). **b**, Cleavage activity of DO1Cas9 and SpCas9, synthesized by in vitro transcription-translation (IVTT), at varying dilutions (10×, 100×, 1000×) on a PAM library. Bands indicate PCR-amplified adapter-ligated cut fragments^8^. **c**, Left: WebLogo of PAM preferences from position frequency matrices (PFMs) of DO1Cas9 (100nM, 100× dilution) cut sites in a plasmid library Right: nucleotide permissiveness within the 7-nucleotide PAM region (green, full; yellow, moderate; orange, low; red, none). **d**, Phylogenetic tree of Cas9 nucleases generated using iTOL^25^ (Interactive Tree of Life) from sequences retrieved from the Caspedia Database^9^ and the Malaspina Dataset^6^, colored by Class 2 type (II-A, II-B, II-C). **e**, DO1Cas9 structure modeled with AlphaFold. **f**, Local structural flexibility (RMSF) for DO1Cas9, SpCas9, and GeoCas9 AlphaFold models. **g**, Predicted free energy of unfolding (kcal/mol) for the same three proteins. **h**, Unfolding dynamics for DO1Cas9 and SpCas9 ± gRNA, by nanoDSF. **i**, Top: pipeline testing how temperature pre-treatment of DO1Cas9 and SpCas9 apo-forms affects subsequent RNP nuclease activity. Bottom: in vitro nuclease activity of DO1Cas9 and SpCas9 RNPs after incubating the apo-forms at different temperatures for 1h. **j**, Top: scheme of capillary electrophoresis-based fragment analysis. Bottom: percentage of DNA cleavage by DO1Cas9 across 15–40°C (mean ± SD, N = 3). **k**, Left: setup for low temperature digestions. Middle: in vitro digestion by DO1Cas9 and SpCas9 RNPs at low (7.6°C, 120 min) and moderate (20°C, 30 min) temperatures. Right: percentage of DNA cleavage by DO1Cas9 and SpCas9 at ∼7.6°C at different timepoints, analyzed by capillary electrophoresis (mean ± s.d.; DO1Cas9, N = 3; SpCas9, N = 2). **l**, In vitro digestion of *gfp* PCR amplicons by DO1Cas9 and SpCas9 using gRNAs with a perfectly matched spacer (M) or with mismatches at positions +1, +3, and +5. C, undigested control.

DO1Cas9 is a class 2 type II-C Cas9 that shares very low sequence identity with the widely used SpCas9 (type II-A) and is also distinct from other type II-C nucleases such as Nme2Cas9 or Cme2Cas9 (**Extended Data Fig. 1**), the latter being the closest evolutionary relative of DO1Cas9 among Cas9 orthologs with an identified PAM (**Fig. 1d**)^9^. Intriguingly, Cme2Cas9 is rather a thermophilic nuclease^2^, in contrast to the psychrophilic origin of DO1Cas9.

Structural models generated by AlphaFold^10^, together with structural and sequence alignments to other Cas9 proteins, identified histidine 574 as a catalytic residue within the HNH domain of DO1Cas9. This residue, together with aspartate 9 in the RuvC domain, was found to be essential for ribonucleoprotein (RNP) nuclease activity by generating the D9A/H574A nuclease-dead variant (**Fig. 1e and Extended Data Fig. 2**). AlphaFold2/ColabFold ensembles of apo DO1Cas9 revealed pronounced conformational heterogeneity, indicating high structural flexibility, a hallmark of enzymatic adaptation to cold environments^11^. Accordingly, DO1Cas9 showed higher root-mean-square fluctuation (RMSF) values than mesophilic SpCas9 and thermophilic GeoCas9 in the HNH catalytic domain and the PAM-interacting region, indicating enhanced mobility in regions directly involved in target recognition and cleavage (**Fig. 1f**). Such increased flexibility at functionally important sites is thought to compensate for the reduced molecular motions at low temperatures, thereby helping to preserve catalytic activity in cold environments ^11^.

Given its marine origin, we examined the salt-dependent stability of DO1Cas9 by calculating the apo-state surface electrostatic potential (ϕ) using APBS^12^ (Adaptive Poisson-Boltzmann Solver) across ionic strengths, with SpCas9 as a non-marine reference (**Extended Data Fig. 3**). DO1Cas9 displayed a highly positive surface potential that promotes electrostatic repulsion and destabilization among protein regions under low-salt conditions. Increasing NaCl to 500mM, a concentration similar to its marine environment, screens this repulsion and restores stability. This prediction was consistent with the salt-dependent thermal stability observed experimentally by nano-Differential Scanning Fluorimetry (nanoDSF) (**Extended Data Fig. 4**).

Since the DO1Cas9 sequence was retrieved from bacteria (Bacteroidetes phylum)^6^ living in the deep ocean at low temperatures, we asked whether it exhibits other features of cold-adapted enzymes^11,13,14^ beyond enhanced structural flexibility. By predicting absolute protein stability using generative models^15^, we found that DO1Cas9 has reduced thermodynamic stability compared with SpCas9 and GeoCas9 (**Fig. 1g**). We also used nanoDSF to monitor heat-induced denaturation from the folded state of the apo forms of DO1Cas9, SpCas9 and the Cryonase cold-active nuclease derived from a psychrophilic bacterium (*Shewanella* sp.). DO1Cas9, similarly to Cryonase, showed a shallow sigmoidal unfolding profile, indicating a gradual denaturation process initiated at low temperatures in contrast to the sharp unfolding transition observed for SpCas9 (**Fig. 1h and Extended Data Fig. 5**). In line with previous studies^16^, gRNA binding increased thermal stability to both Cas9 nucleases (**Fig. 1h**).

To study the low thermal stability of the DO1Cas9 apo-enzyme we pre-incubated the protein at different temperatures before assessing RNP activity. Unlike SpCas9, a 15-min incubation at 33°C under standard salt concentration (100mM NaCl) was sufficient to inactivate DO1Cas9’s nuclease activity. This property enables selective thermal denaturation of DO1Cas9 during biotechnological processes (**Fig. 1i and Extended Data Fig. 6**). As an RNP, DO1Cas9 activity decayed sharply beyond 35°C (**Fig. 1j**), unlike mesophilic nucleases, which remain active over a broader temperature range^2^.

To directly test whether DO1Cas9 retains catalytic activity under conditions that would inactivate a mesophilic nuclease, we performed in vitro cleavage assays under tightly controlled cold conditions. DO1Cas9 cleaved dsDNA at 7.6°C within a 30-min reaction, whereas SpCas9 did not (**Fig. 1k**). Using capillary electrophoresis for quantification, SpCas9 showed low activity only after extended incubation, reaching roughly a third of DO1Cas9’s cleavage (**Fig. 1k**). As an additional distinctive feature, DO1Cas9 showed greater sensitivity than SpCas9 to guide-target mismatches, suggesting high target specificity (**Fig. 1l and Extended Data Fig. 7**).

We investigated whether the peculiar DO1Cas9 structural and biophysical properties would limit its nuclease activity in different organisms, starting with *Caenorhabditis elegans*, which is an amenable animal model to evaluate CRISPR-Cas activities^17^. DO1Cas9 RNPs targeting *dpy-10* were microinjected into the germline of N2 hermaphrodites with fluorescent co-injection markers. DO1Cas9 efficiently edited the *dpy-10* locus using distinct gRNA formats (sgRNA (commercial of IVT), or dual crRNA:tracrRNA) across a range of temperatures (10, 15 and 20 °C), with 30% to 50% of fluorescent F1 animals segregating dumpy F2 progeny (**Fig. 2a and Extended Data Fig. 8**). To assess RNP persistence in vivo, we engineered a fluorescent reporter that recovers fluorescence upon Cas9-mediated non-homologous end joining (NHEJ) repair. Sparse red fluorescent cells appeared in F1 somatic tissue as a result of mosaicism^18^ generated by Cas9 activity in embryonic or post-embryonic somatic cells. This indicates that injected RNP remains active for at least several hours after germline injection, persisting from progenitor germ cells to somatic cells in the progeny (**Fig. 2b**).

**Figure 2.**
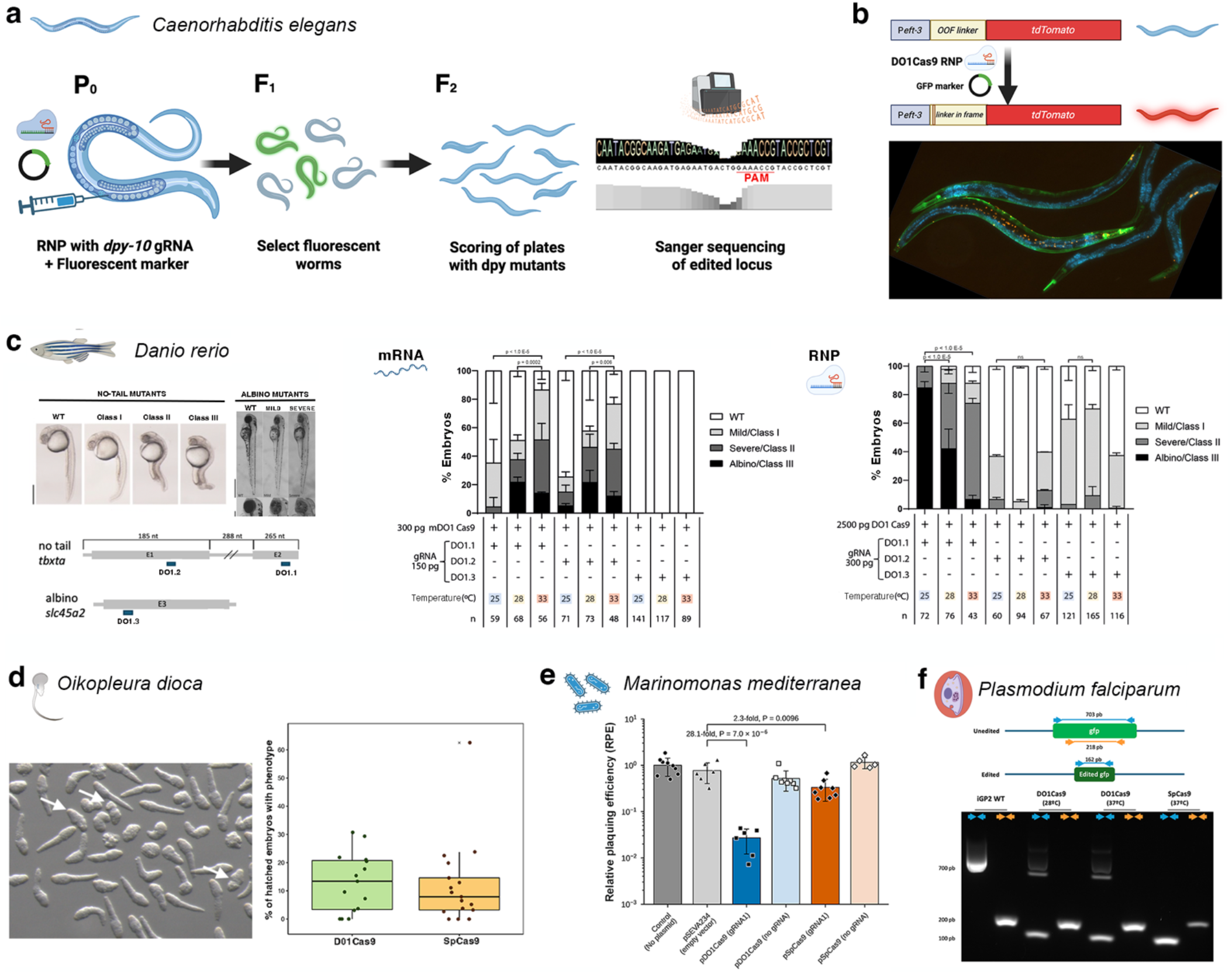
DO1Cas9 activity in vivo. **a**, Strategy for gene editing with DO1Cas9 in *C. elegans*. DO1Cas9 RNPs and a fluorescent co-injection marker were injected into the germline. Nuclease activity produces dumpy phenotype in F2 generation. Sanger sequencing confirms mutations at the PAM-proximal *dpy-10* target. **b**, Top: tdTomato sensor construct integrated in the *C. elegans* genome. Nuclease activity at the out-of-frame linker would restore the reading frame and the fluorescence. Bottom: example of worms with restored red fluorescence in somatic cells. Green fluorescence (P*mlc-1*::gfp) marks successful microinjection. **c**, DO1Cas9 gene editing in zebrafish. Left: no tail and albino phenotypes at 28 and 48 hours post hatching (hpf), respectively. Dark blue rectangles indicate targeted regions in *tbxta* and *slc45a2* genes. Middle/Right: editing efficiencies at *tbxta* and *slc45a2* after injecting DO1Cas9 mRNA or RNP, across three temperatures (n = embryos analyzed; ≥2 independent experiments, χ^2^ test, p ≥ 0.05). **d**, *O. dioica* embryos injected with either DO1Cas9 or SpCas9 targeting *Brachyury*. Left: representative “shrunken tail” phenotype (arrows), absent in control clutches. Right: boxplot showing the proportion of embryos exhibiting the Bra phenotype among total hatched embryos. Because females were injected, spawned, and fertilized individually, each clutch was considered an independent biological replicate. Dots represent individual clutches. No significant difference was detected between Cas9 nucleases (Mann-Whitney U test, *p* = 0.557). **e**, Protective effect of DO1Cas9 and SpCas9 guided by a gRNA against phage CB5A infection in *M. mediterranea* bacteria lacking native CRISPR systems. Bars indicate the mean and error bars the standard deviation; each symbol represents one independent replicate. Statistical significance was assessed by two-sided unpaired Welch’s t-tests on log_10_-transformed RPE values, comparing pDO1Cas9 (gRNA1) and pSpCas9 (gRNA1) against the empty vector (pSEVA234); fold-reduction in mean RPE relative to the empty vector is indicated above each comparison. **f**, DO1Cas9 gene editing by HDR in *P. falciparum*. Cas9 plasmids were electroporated and transformed parasites were drug-selected at 37°C and incubated at either 37°C or 28°C for 36 h. HDR-mediated shortening of the gfp locus was assessed by PCR.

In zebrafish, we used DO1Cas9 to target three protospacers in two genes: one involved in the formation of the posterior part of the embryo (*tbxta*) and one required for pigmentation (*slc45a2*), at 25, 28 and 33ºC. With DO1Cas9 mRNA, no-tail phenotype penetrance increased with temperature at *tbxta* sites, likely reflecting improved mRNA translation at higher temperatures (**Fig. 2c**). In contrast, the penetrance of phenotypes using DO1Cas9 RNP was significantly higher at lower temperatures (**Fig. 2c**).

The marine nature and salt requirements would make the DO1Cas9 suitable for applications in marine species. We tested DO1Cas9 and SpCas9 in the planktonic animal *Oikopleura dioica* by injecting the RNPs into the premature female gonad targeting the *Brachyury* gene and they showed comparable editing efficiency, producing the expected “shrunken tail” phenotype^19^ with notochord defects in ∼13% of embryos (**Fig. 2d**). We also assessed DO1Cas9 in *Marinomonas mediterranea* MMB-1, a marine Gram-negative bacterium that defends against podovirus infections via coordinated activity of the resident I-F and III-B CRISPR-Cas systems targeting an early phage gene (gp6)^20^. In laboratory conditions, upon viral infections, *M. mediterranea* CRISPR-Cas defense system reduces the number of viral Plaque Forming Units (PFU)^20^. In *M. mediterranea* ΔI-F ΔIII-B, a strain lacking both endogenous CRISPR systems, plasmids expressing DO1Cas9 and SpCas9 with a gp6-targeting sgRNA conferred phage resistance against phage CB5A (**Fig. 2e**). Although both pDO1Cas9 and pSpCas9 interfered with the phage infection, the magnitude of interference differed substantially: pDO1Cas9 reduced the Relative Plaquing Efficiency (RPE) by 28-fold, consistent with efficient phage interference, whereas pSpCas9 reduced RPE by only 2.3-fold, a modest effect of limited practical antiviral value.

Finally, we assessed DO1Cas9 editing by homologous recombination (HR) in *Plasmodium falciparum*, which lacks the canonical NHEJ pathway (**Extended Data Fig. 9**). At both 28°C and 37°C, double cuts generated by DO1Cas9 were repaired by homology-directed repair (HDR) using a donor DNA template (**Fig. 2f**).

Although editing efficiencies could likely be further improved through reaction optimization or protein engineering, our results establish DO1Cas9 as a cold-active genome-editing nuclease functional across diverse organisms. Recently, a cold-adapted Cas9 (Fp2Cas9) from the fresh-water bacterium *Flavobacterium psychrophilum* has been described^21^. Unlike Fp2Cas9, DO1Cas9 is about one-third smaller (991 vs. 1548 aa) and originates from a salty, deep-water environment, which confers distinct biophysical properties and requirements on DO1Cas9. Moreover, we demonstrated that, under tightly controlled conditions, DO1Cas9 cuts dsDNA at low temperatures where mesophilic SpCas9 cannot. Therefore, its potential for editing genes under specific and exclusive cold and salt conditions sets DO1Cas9 as a versatile addition to the Cas9 toolbox for biotechnological applications^22,23^, such as controlled activation switches in molecular diagnostics or genome editing under cold-chain storage conditions.

## SUPPLEMENTARY DATA

**Supplementary Data 1:** DO1Cas9 DNA, protein, crRNA and tracrRNA sequences. DNA and predicted protein sequences of DO1Cas9 identified from deep-ocean metagenomic data. crRNA and tracrRNA sequences detected in the DO1Cas9 CRISPR-Cas locus and used for sgRNA design in this study.

**Supplementary Data 2:** PAM determination: plasmid library and in vitro experiments. Sheets 1-5 contain data from the IVTT-based PAM determination assay performed with 100nM DO1Cas9, including the position frequency matrix (PFM), PAM enrichment analysis, top PAM counts, PAM-distal sequence analysis, and cleavage position summary. Sheet 6 lists the guide RNAs used for cleavage assays on PCR templates containing different PAM sequences.

**Supplementary Data 3:** gRNA sequences, primers and plasmids used in this study.

**Supplementary Data 4:** Cas9 sequences for phylogenetic analyses from CasPEDIA database and Malaspina expedition dataset. List of 48 sequences from Malaspina expedition dataset and 830 sequences retrieved from CasPEDIA database.

**Supplementary Data 5:** Uncropped gel images

## EXTENDED DATA

**Extended Data Fig. 1** Alignment of protein sequences for DO1Cas9, SpCas9, Nme2Cas9 and Cme2Cas9

**Extended Data Fig. 2** Validation of DO1Cas9 dead variant

**Extended Data Fig. 3** Surface electrostatic properties of DO1Cas9 and SpCas9

**Extended Data Fig. 4** Influence of salt concentration in the thermal stability of apo DO1Cas9

**Extended Data Fig. 5** DO1Cas9, SpCas9 and Cryonase thermal-response profiles at nanoDSF

**Extended Data Fig. 6** Thermal instability of DO1Cas9 apo-form at 33°C

**Extended Data Fig. 7** DO1Cas9 sensitivity to gRNA-target mismatches (*dpy-10* target)

**Extended Data Fig. 8** DO1Cas9 gene editing in *C. elegans*: **a**, CRISPR mixes and efficiencies. **b**, Sanger sequencing of representative edits.

**Extended Data Fig. 9**. Schematic of the strategy for HDR-mediated genome editing of the *gfp* locus in *Plasmodium falciparum* using DO1Cas9 or SpCas9.

## ACKNOWLEDGEMENTS

This work was mainly supported by a Proof-of-concept grant, reference PDC2021-120968-I00, to SGA, and a *Proyecto General del Conocimiento* grant, reference PID2023-146930NB-I00 to JC, both from the Spanish Ministry of Science and Innovation. AOL has a predoctoral FPI contract, reference PREP2023-001127. F.J.M.M. is supported by grants PID2023-150750NB-I00 (funded by MICIU/AEI/ 10.13039/501100011033 and by ERDF/EU), and INNEST/2024/427 (funded by Agencia Valenciana de Innovación - IVACE+I Innovación – and by the European Union through the ERDF Program of the Valencian Community 2021-2027). AG was supported by the Ministerio de Ciencia e Innovación, which is part of Agencia Estatal de Investigación (AEI), through the Retos Investigación Grant PID2022-140457OB-I00 funded by MCIU /AEI /10.13039/501100011033, grant no. AC23_2/00040 by ISCIII through AES 2023 and within the European Joint Programme Rare Diseases framework, by Instituto de Salud Carlos III (ISCIII), and “Programa FORTALECE del Ministerio de Ciencia e Innovación” through the project number FORT23/00032. D.R.M. was supported by the Scientific Foundation of the Spanish Association Against Cancer project number INVES2610039ROME. M.M.M. was supported by the Government of Andorra with the predoctoral fellowship, ATC030-AND/2022. We thank Till Voss for the NF54-iGP2 parasite line and Abdouramane Traore for helping with transgenic parasite cloning. EGD was supported by the La Caixa Foundation-Health Research 2020 Call [HR20-00635 to E.G.D]. RSD was funded by FPU fellowship (grant no. FPU20/04280) of the Spanish Ministry of Universities. CrC was supported by the Ministerio de Ciencia, Innovación y Universidades, Gobierno de España [grant number PID2025-170547NB-I00 and PID2022-141627NB-I00], the Institució Catalana d’Investigació i Estudis Avançats Acadèmia (ICREA), Generalitat de Catalunya [grant number Ac2215698]; and Agència d’Ajuts Universitaris i de Recerca, Generalitat de Catalunya [grant number 2021-SGR00372]. MAMM lab has been supported by Grant PID2024-158755NB-I00 funded by MICIU/AEI/10.13039/501100011033 and FEDER, UE and co-financed by the Spanish Ministry of Science and Innovation with funds from the European Union NextGenerationEU (PRTR-C17.I1) and the Regional Ministry of University, Research and Innovation of the Autonomous Community of Andalusia within the framework of the Biotechnology Plan applied to Health. CAM has a predoctoral contract by the Ministerio de Ciencia, Innovación y Universidades, Gobierno de España [FPU23/02391 and FPI:PREP2022-000800]. A M-C was supported by pre-doctoral contract by the University of Murcia. We thank Felipe Coutinho for sharing sequences and bioinformatic tools.

## AUTHOR CONTRIBUTIONS

SGA and JC envisioned the original idea of this project, secured funding to support this study and coordinated the collaborative network (Deep Cas consortium). AOL performed most of in vitro experiments and *C. elegans* assays. SGA, AOL and JC managed scientific and logistic interactions with collaborators. SGA, PS and AB worked on metagenomes and led bioinformatic analyses. JC contributed to in vivo *C. elegans* experiments. MS, TU and GG contributed to determine DO1Cas9 PAM sequence. FC performed AlphaFold analyses. GM, MM and DR contributed at early stages of the project as master’s students. GM analyzed DO1Cas9 homologies and identities with other Cas9s. CSB and MAMM led and performed zebrafish experiments. ASA and AMC led and performed *M. mediterranea* experiments. CAM, AFR and Cr.C led and performed *Oikopleura* experiments. RSD, DLF, and EGD led and performed *P. falciparum* experiments. JG contributed with biophysical studies. FM helped in the initial sequence analyses from metagenomes and kept a continues activity as advisor through the project. MMM, DRM, AG and AOL were involved in exploratory experiments with cell lines. NRM, CC, DCO, AVM and SRM were involved in DO1Cas9 protein purifications. JC and AOL wrote the manuscript and assembled figures for publication.

## COMPETING INTERESTS

The authors declare the following competing interests: T.U. and G.G. are employees of CasZyme, a company developing and commercializing CRISPR-Cas-based genome-editing tools. M.S: was an employee of CasZyme at the time of the study. S.G.A. and J.C. are co-inventors on a pending patent application related to DO1Cas9. The remaining authors declare no competing interests.

### Animal Ethics declaration

not applicable

### Data availability

The data used for generating the figures are found in Supplementary Data 1–5. All other data or materials supporting the points of this study are available from the corresponding author upon reasonable request.

## METHODS

### Identification of the nuclease DO1Cas9 and its guide RNA

The DO1Cas9 nuclease was identified from metagenomic datasets generated during the Malaspina Expedition 2010, a global oceanographic circumnavigation conducted aboard the R/V Hespérides. Samples were collected from 33 oceanographic stations, including 13 vertical profiles spanning depths from the surface to 4,000 m. DO1Cas9 originated from a metagenomic sample collected at a depth of 1,500 m in the Atlantic Ocean (station 135). A non-redundant gene catalog comprising 17.4 million predicted genes from the Malaspina metagenomes was screened for CRISPR-Cas systems, yielding 48 Cas9 proteins^6^. Seven Class 2 type II Cas9 candidates were selected for experimental validation, and DO1Cas9 was among the nucleases exhibiting high DNA-cleavage activity in a plasmid library assay.

The DO1Cas9 CRISPR-Cas locus was analyzed using PILER-CR v1.06, CRISPRCasFinder v1.1.2, and Prokka v1.14.6 to identify the associated guide RNA. Putative tracrRNA sequences were inferred through a multi-step computational workflow. First, CRISPR repeat sequences were aligned against the corresponding locus-containing contig using BLASTN (v2.7.1; parameters: -blastn-short -dust no). Sequences located within 500 bp upstream or downstream of the CRISPR-Cas locus and displaying similarity to the CRISPR repeat were considered candidate antirepeat regions. Second, Rho-independent-like termination signals were identified in the vicinity of candidate antirepeats using the FindTerm web server. Third, putative promoter sequences were predicted using BPROM. Finally, interactions between candidate antirepeats and CRISPR repeats were evaluated using IntaRNA v3.3.1.

Candidate tracrRNAs were defined as sequences extending from the antirepeat to the downstream Rho-independent termination signal, including the poly(U) tract, when all of the above criteria were satisfied. The transcriptional orientation of the CRISPR array was inferred from the highest-scoring repeat-antirepeat interactions and the predicted tracrRNA organization. All coordinates are reported relative to the original contig sequence in the positive orientation.

### RNP activity on plasmids and Identification of the PAM sequence

DO1Cas9 ribonucleoprotein (RNP) complexes were generated using an *E. coli* cell-free in vitro transcription-translation (IVTT) system as previously described^26^. sgRNA was transcribed directly in the IVTT reaction from a double-stranded DNA template containing a T7 promoter and the sgRNA coding sequence. Briefly, 250 ng of plasmid encoding DO1Cas9 and a 100-fold molar excess of sgRNA template were added to the IVTT reaction mixture. Subsequently, 5μl of the resulting Cas9-sgRNA complex, or serial tenfold dilutions thereof, were incubated with 0.5μg of a previously described plasmid library containing a randomized 7-bp PAM sequence^8^ in a 50μl reaction consisting of 10mM Tris-HCl (pH 7.5), 100mM NaCl, 10mM MgCl2, and 1mM DTT. Reactions were incubated for 60min at 37 °C.

Cleaved library fragments were enriched by adapter ligation and PCR amplification prior to next-generation sequencing. DNA ends were repaired by incubation with T4 DNA polymerase (1 U; New England Biolabs, M0203S) and dNTPs (0.3μl of a 10mM mix; Thermo Fisher Scientific, R0192) for 15min at 12 °C, followed by heat inactivation at 75°C for 20 min. A 3^′^ dA overhang was subsequently added using DreamTaq DNA polymerase (1.5 U; Thermo Fisher Scientific) for 30 min at 72 °C. DNA was purified using Monarch PCR & DNA Cleanup columns (New England Biolabs) and ligated to adapters carrying a complementary 3^′^ dT overhang using T4 DNA ligase (400 U; New England Biolabs) in 25μl of ligation buffer containing 50mM Tris-HCl (pH 7.5), 10mM MgCl2, 10mM DTT, 1 mM ATP, and 5% (w/v) PEG 4000. Following incubation for 1 h at room temperature, 10μl of the ligation reaction was amplified using Q5 DNA polymerase (New England Biolabs; 15 cycles) with primers specific to the PAM-proximal region of the library and the ligated adapter (Supplementary Data 3). PCR products were purified, and Illumina sequencing adapters and sample indices were incorporated through two successive rounds of PCR using Phusion High-Fidelity PCR Master Mix in HF buffer (New England Biolabs; 10 cycles per round). Libraries were sequenced on an Illumina MiSeq instrument with a 25% (v/v) PhiX Control v3 spike-in.

PAM sequences supporting double-stranded DNA cleavage were identified as previously described^2^. Following sequencing, cleavage sites within the protospacer were determined by identifying the position with the highest abundance of adapter-ligated reads using a custom analysis script. PAM sequences associated with cleavage events were extracted and their nucleotide frequencies at each randomized position were compared with those of the input library using the normalization formula (treatment frequency)/((control frequency)/(average control frequency)). PAM preferences were quantified as position frequency matrices (PFMs) and visualized using WebLogo. To minimize background arising from non-specific cleavage associated with the IVTT reaction components, analyses were restricted to the top 10% most abundant PAM sequences.

### Phylogenetic analyses

To generate a phylogenetic tree, the 48 Cas9 sequences obtained from the Malaspina dataset^6^ were combined with 830 Cas9 sequences from the CasPEDIA database^9^ (Supplementary Data 4) and aligned with Muscle v5.161 using the “-super5” parameter. Phylogenetic reconstruction was performed through IQ-TREE v2.0.662 using the “-MFP” option, which identified VT+F+R10 as the best substitution model. The generated tree was visualized using iTOL21.

### 3D-protein models, RMSF and stability predictions

Structural models of DO1Cas9 in complex with guide RNA and target DNA were generated using the AlphaFold3 server with default parameters^10^. The resulting models were used for structural analyses. AlphaFold3 prediction files have been deposited at https://github.com/fcolizzi/DO1_Cas9.

To assess conformational dynamics and thermal stability, conformational ensembles of apo Cas9 proteins (DO1Cas9, SpCas9 (UniProt: Q99ZW2), and GeoCas9 (UniProt: A0A250DVH8)) were generated using the ColabFold implementation of AlphaFold2^15,27^. Structural ensembles were derived from a single input sequence by increasing uncertainty within the AlphaFold2 prediction network. Two complementary sampling strategies were employed: (i) reduction of multiple sequence alignment (MSA) depth (max_msa = 32:64, num_seeds = 16)^28^ and (ii) activation of dropout layers during inference (use_dropout, num_seeds = 16)^29^. Predictions generated using both approaches were combined to obtain an ensemble of conformations for each protein. Root mean square fluctuation (RMSF) and thermal stability analyses were subsequently performed on the resulting ensembles^30^.

### Protein purification

DO1Cas9 carrying a C-terminal SV40 nuclear localization sequence and His_6_ tag (Supplementary Data 3) was expressed in *Escherichia coli* BL21(DE3) pLysS cells from plasmid pTEM14_ProteaseIV. Expression was induced with 0.2mM IPTG at an OD_600_ of 0.6-0.8, and cultures were incubated overnight at 16°C. Cell pellets were resuspended in 50 mL Buffer A (50mM TRIS HCl ph7.4, 500mM de NaCl, 10mM imidazole, 2mM DTT, 10% glycerol) supplemented with EDTA-free protease inhibitor cocktail (Merck, 4693116001). Cells were lysed by French press and the soluble fraction was purified by Ni^2+^ affinity chromatography followed by size-exclusion chromatography on a Superdex 200 16/600 column equilibrated in 40mM Tris-HCl (pH 7.4), 1M NaCl, 0.2 mM EDTA, and 2 mM DTT. Purified protein was exchanged into storage buffer containing 20mM Tris-HCl (pH 7.4), 500mM NaCl, 1mM DTT, 0.1mM EDTA, and 50% glycerol and stored at −20°C. Protein purity was assessed by SDS-PAGE.

### sgRNA production by in vitro transcription

Single-guide RNAs (sgRNAs) were synthesized by in vitro transcription (IVT) using the HiScribe^®^ T7 Quick High Yield RNA Synthesis Kit (NEB, #E2050). Briefly, DNA templates containing a T7 promoter upstream of the sgRNA sequence were obtained either as gBlocks^®^ Gene Fragments (Integrated DNA Technologies, IDT) or as gene fragments (GenScript) and were used as substrates for T7 RNA polymerase-mediated transcription (Supplementary Data 3). Transcription reactions were incubated according to the manufacturer’s recommendations, after which template DNA was removed by DNase I treatment.

The resulting sgRNAs were purified using the Monarch^®^ RNA Cleanup Kit (NEB, #T2050). Purified sgRNAs were eluted in nuclease-free water, quantified by spectrophotometry, and stored at −80°C until use.

### NanoDSF analyses of protein stability

Protein thermostability was assessed by nano differential scanning fluorimetry (nanoDSF)^31^ using a Prometheus NT.48 instrument (NanoTemper Technologies). Thermal unfolding was monitored by measuring intrinsic tryptophan and tyrosine fluorescence at 330 and 350 nm, and the melting temperature (T_m) was determined from the temperature-dependent change in the F330/F350 fluorescence ratio. T_m was defined as the temperature corresponding to the midpoint of the unfolding transition^32^.

Protein samples were prepared at 1mg/ml in the indicated buffers, and 10μl of each sample was loaded into standard capillaries. Prior to analysis, samples were centrifuged at 16,000 × g for 10 min at 4°C to remove insoluble material and protein aggregates. Excitation power was adjusted to obtain fluorescence intensities exceeding 2,000 arbitrary units at both wavelengths. Samples were heated from 20°C to 95°C at a rate of 1°C/min, and unfolding transitions were analysed using the manufacturer’s software.

### Electrostatic potential calculations

Electrostatic potentials were computed by solving the linearized Poisson–Boltzmann equation with APBS^12^ (v3.4.1). Atomic partial charges and radii were assigned with PDB2PQR^33^ using the AMBER force field, with protonation states set at pH 7.0 as determined by PROPKA. Calculations used a solute dielectric of 2.0, a solvent dielectric of 78.54, a solvent probe radius of 1.4 Å, a temperature of 298.15 K, and grid spacing derived automatically from the molecular dimensions. Monovalent ion species (radius 1.62 Å) were included explicitly to model ionic strength; potentials were computed at 0.1 M and 0.5 M NaCl to bracket the standard assay (100mM) and seawater/high-salt (500mM) conditions. Three-dimensional models of DO1Cas9 and SpCas9 were generated with AlphaFold3^10^. Apo-state models were used for all electrostatic analyses.

Surface electrostatic metrics were derived by sampling the potential on a shell 2–5 Å from the solvent-accessible surface. We report the mean surface potential (ϕ, in kT/e) and the positive-and negative-patch areas, defined as the fraction of the sampled surface with ϕ > +1 kT/e and ϕ < −1 kT/e, respectively. All proteins were processed with identical parameters to ensure comparability. Dipole moments and net charges were computed from the PQR atomic charges.

### In vitro DNA cleavage assays

Guide RNAs were generated by in vitro transcription from synthetic double-stranded DNA templates (gBlocks,IDT; gene fragments, GenScript) using the HiScribe Quick T7 High Yield RNA Synthesis Kit (NEB, E2050). Unless otherwise indicated, ribonucleoprotein (RNP) complexes were assembled by incubating purified Cas9 protein (350 nM) and sgRNA (350 nM) in 1× NEBuffer r3.1 (NEB, B7203S) in a final volume of 30 μl for 30 min at 20 °C.

DNA cleavage reactions were initiated by the addition of a PCR-amplified fragment of the *dpy-10* or *gfp* genes (final concentration, 200 nM). Sequences of primers for DNA substrates and guide RNAs are provided in Supplementary Data 3. Unless otherwise specified, reactions were incubated for 30 min at 25 °C. Reactions were terminated by the addition of 6 μl BlueJuice loading dye containing EDTA (Invitrogen, 10666143) and 1 μl Proteinase K (20 mg ml^−1^), followed by incubation at 56°C for 10 min. Unless otherwise indicated, cleavage assays were performed in a final volume of 30 μl and were resolved by electrophoresis on 1.5% agarose gels stained with SYBR™ Safe DNA Gel Stain (Invitrogen, S33102). Assays intended for capillary electrophoresis analysis were performed in a final volume of 20 μl and processed as described below.

#### Protein stability assays

To evaluate protein stability, purified Cas9 protein was incubated in 1× NEBuffer r3.1 at the indicated temperatures for 1 h before RNP assembly and DNA cleavage assays. For time-course experiments at 33°C, proteins were pre-incubated for 5, 15, 30, or 60min before RNP assembly and subsequent cleavage assays.

#### Temperature-dependent activity

For temperature-dependent activity measurements, RNP complexes were assembled using 200nM Cas9 protein and 400 nM sgRNA in a final volume of 20 μl, as described above. Cleavage reactions were performed at 15, 26, 30, 35, 37, or 40°C for 30min. Cleavage products were quantified by capillary electrophoresis as described below.

#### Sensitivity to mismatches

To assess mismatch tolerance, RNP complexes were assembled using in vitro-transcribed guide RNAs that were either fully complementary to the target sequence or contained single mismatches within the spacer region at positions +1, +3, or +5 relative to the PAM-proximal end of the spacer. DNA cleavage assays were subsequently performed under the standard reaction conditions described above.

#### Low-temperature cleavage assays

Low-temperature cleavage assays were performed under strictly controlled conditions to minimize temperature fluctuations. After assembly of RNPs, the complexes and DNA substrates were pre-cooled separately for 10 min before reaction initiation. Cleavage reactions were initiated by addition of the DNA substrate and incubated at 7.6°C for the indicated time periods using a thermoblock (PCH-1, Grant Bio). The setup was kept inside a 15°C incubator to maintain a controlled environmental temperature. Reaction temperature was monitored using a dual-channel thermocouple thermometer (PST262, Proster). Reactions were terminated and analysed either by agarose gel electrophoresis or capillary electrophoresis.

#### Capillary electrophoresis analysis of cleavage products

Fluorescently labelled double-stranded DNA substrates were generated by PCR amplification using a 5^′^-6-FAM-labelled oligonucleotide (Integrated DNA Technologies). Cleavage reactions were performed under the standard conditions described above, except that reactions were quenched by addition of 0.5 M EDTA to a final concentration of 25 mM, followed by addition of 1 μl Proteinase K (20 mg ml^−1^) and incubation at 56 °C for 10 min to deproteinize the reaction mixture. Then, reaction was incubated at 95° C for 10 min to inactivate Proteinase K. Samples were subjected to fragment analysis by capillary electrophoresis (Macrogen). Electropherograms were analysed using Peak Scanner 2 software (Applied Biosystems). The percentage of digested product was calculated from the Area-in-Point peak area metric as:

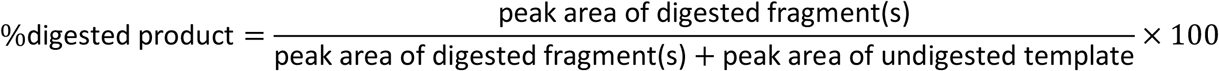

Data are presented as mean ± standard deviation (SD). Temperature-dependent DO1Cas9 activity was determined from three independent measurements (n = 3). For the low-temperature (∼7.6°C) time-course assay, DO1Cas9 activity was determined from three independent measurements (n = 3), whereas SpCas9 activity was determined from two independent measurements (n = 2). No inferential statistical analyses were performed.

### Genome editing in *Caenorhabditis elegans*

#### Microinjection procedure

CRISPR injection mixes were prepared by combining equimolar concentrations of Cas9 nuclease, tracrRNA, and crRNA. DO1Cas9 RNP complexes were assembled by incubation at 20°C for 30 min. Injection mixes were centrifuged at 16,100 × g for 2 min and kept on ice until use. Young adult hermaphrodites were immobilized on 2% agarose pads under halocarbon oil and microinjected using a XenoWorks Microinjection System (Sutter Instrument) coupled to a FemtoJet microinjector (Eppendorf) and a Nikon Eclipse Ti-S microscope equipped with Nomarski optics. Approximately 15-20 animals were injected for each experimental condition. Following injection, animals were recovered in M9 buffer and transferred individually to nematode growth medium (NGM) agar plates^34^ and allowed to produce progeny.

#### CRISPR-Cas editing of dpy-10

N2 (VC2010 reference assembly) hermaphrodites were injected with DO1Cas9 ribonucleoprotein complexes targeting the *dpy-10* locus together with fluorescent co-injection marker plasmids (Extended Data Figure 8). Following microinjection, animals were recovered on individual NGM plates and allowed to produce progeny. Fluorescent F1 animals were isolated individually and transferred to fresh plates. Genome editing activity was assessed by screening the F2 progeny of fluorescent F1 animals for the Dumpy (*Dpy*) phenotype, indicative of mutagenesis at the *dpy-10* locus. Editing efficiency was calculated as the proportion of fluorescent F1 animals that produced Dumpy animals in the F2 progeny. For molecular characterization of editing outcomes, genomic DNA was prepared from independent mutant isolates, and the targeted *dpy-10* locus was amplified by PCR and analysed by Sanger sequencing. Sequence variants were identified by comparison with the reference *dpy-10* sequence.

#### In vivo fluorescent reporter assay

The Cas9 activity reporter strain CER780 was generated by CRISPR-Cas9-mediated insertion of an out-of-frame (+1 frameshitf) linker into the EG7902 reporter strain, originally developed by the Frøkjær-Jensen laboratory^32^. The linker sequence was 5’-TCCTCCGGAAATTCCAATGCCAAATTCCCGGGGACCATCCTTTTCCTCCGGACTTGTTCCACTTTCCCTTAGGGGATCCCAT-3’. EG7902 carries a P*eft-3*::tdTomato reporter cassette, and insertion of the linker between the P*eft-3* promoter and the tdTomato coding sequence disrupts the reading frame, resulting in loss of tdTomato fluorescence. Genome editing events (indels) that restored the correct reading frame led to recovery of red fluorescence.

To assess DO1Cas9 activity, CER780 animals were microinjected with a mixture containing DO1Cas9 RNP complexes assembled at a final concentration of 5μM and 260ng P*mlt-2*::GFP plasmid as an injection marker. Injected animals were maintained at 22°C throughout the experiment. GFP-positive F1 progeny were isolated individually and allowed to produce F2 offspring. Editing activity was assessed by quantifying restoration of tdTomato fluorescence in the F2 generation, and representative images were acquired 72 h after injection using a ZEISS Apotome fluorescence microscope.

### Genome editing in zebrafish

#### Preparation of DO1Cas9 mRNA and guide RNAs

A zebrafish codon-optimized DO1Cas9 coding sequence was synthesized (GenScript), cloned into the pT3TS-MCS vector (Addgene #31830), and used for in vitro transcription of capped DO1Cas9 mRNA using the mMESSAGE mMACHINE™ T3 kit (Invitrogen, Thermo Fisher). Following DNase treatment and purification, mRNA was quantified and stored at −80°C. Guide RNAs were generated as described by Vejnar *et al* ^35^ and stored at −80°C until use.

#### Embryo microinjections and temperature treatments

Wild-type AB/Tübingen (AB/Tu) zebrafish were maintained under standard conditions. All animal procedures were conducted in accordance with national and European regulations and were approved by the relevant institutional and governmental ethics committees. To assess DO1Cas9 mRNA activity, 1nl containing 150pg of DO1Cas9 mRNA and 150pg of gRNA was injected into one-cell-stage embryos. For ribonucleoprotein (RNP) delivery, DO1Cas9 protein and gRNA were mixed at a 3.5:1 molar ratio in 25 mM HEPES-KOH (pH 7.5), 250 mM KCl, and 40% (v/v) glycerol, incubated at 25°C for 10 min, and kept on ice prior to injection. One nanoliter of RNP complex (21 fmol) was injected into one-cell-stage embryos.

Following injection, embryos were maintained at 25°C, 28°C, or 33°C for the first 24 h post-fertilization (hpf). Embryos incubated at 25°C or 33°C were subsequently transferred to 28°C, whereas embryos maintained at 28°C remained at that temperature throughout the experiment. Editing efficiency was assessed by scoring tbxta phenotypes at 28 hpf and slc45a2 phenotypes at 48 hpf^36^.

### Genome editing in *Oikopleura dioica*

*Oikopleura dioica* specimens were collected from the Barcelona coastline and have been maintained in laboratory culture for more than 5 years^37,38^. RNP complexes were assembled using either SpCas9 and Alt-R™ sgRNA (IDT) or DO1Cas9 and an in vitro-transcribed sgRNA targeting the Brachyury (Bra) gene (Supplementary Data 3). Cas9 and sgRNA were mixed at equimolar ratios and incubated at 20°C for 20min to a final concentration of approximately 4μM. Phenol red (0.08%) was added to facilitate visualization during injection. Anesthetized (MS222, Sigma-Aldrich) 4-day-old premature female *O. dioica* were microinjected into the ovary using borosilicate needles connected to an InjectMan 4 micromanipulator and a FemtoJet microinjector (Eppendorf). Following injection, animals were maintained overnight at 13°C in seawater supplemented with algae. The next morning, females were individually isolated until spawning, and egg clutches were fertilized with wild-type sperm. Embryos were incubated at 19°C and imaged at 4.5 h post-fertilization (hpf) using a stereomicroscope. Editing activity was assessed by scoring the characteristic *Bra* loss-of-function “shrunken tail” phenotype, characterized by notochord defects and impaired tail elongation. For each clutch, the percentage of hatched embryos displaying the phenotype was calculated.

### Phage resistance assays in *Marinomonas mediterranea*

The DO1Cas9 and SpCas9 coding sequences were codon-optimized for *M. mediterranea* and cloned under the control of the inducible Ptrc promoter in the conjugative vector pSEVA234^39^, generating plasmids pDO1 and pSpCas9. Guide RNAs targeting gene 6 of Murciavirus CB5A (CB5A gp6; Supplementary Data 3) were cloned, in the reverse direction and separated by a transcriptional terminator signal, into pDO1 and pSpCas9 under the constitutive promoter BBa_J23119 (https://parts.igem.org/Part:BBa_J23119). A control plasmid lacking a targeting guide RNA was generated by replacing the guide RNA cassette with an unrelated DNA sequence derived from *trp-lacZ*^*40*^ . Plasmids were synthesized and assembled by Twist Bioscience and mobilized into *M. mediterranea* ΔI-F ΔIII-B by conjugation as previously described^41^. This strain was rendered sensitive to phage CB5A through deletion of its native CRISPR-Cas systems^20^. Cultures were maintained at the standard growth temperature of 25°C, challenged with phage CB5A, and phage resistance was assessed using the double-layer agar assay as previously described^20^. Resistance was quantified as relative plaquing efficiency (RPE), calculated by normalizing plaque counts to those obtained with the parental strain lacking plasmid. Protection mediated by DO1Cas9 and SpCas9 was determined by comparing strains carrying the CB5A gp6-targeting guide RNAs with strains carrying the empty vector or a non-targeting control construct.

### HDR-mediated genome editing in *Plasmodium falciparum*

Parasite line NF54-iGP2^42^ (obtained from Till Voss) was maintained under standard culture conditions in RPMI medium supplemented with 0.5% Albumax II and 2.5mM D-(+)-glucosamine hydrochloride at 3% haematocrit. A DO1Cas9 expression plasmid carrying a gRNA targeting the *gfp* gene and a homologous repair template consisting of a truncated *gfp* fragment (**Supplementary Data 3 and Extended Data Fig. 9**) was generated from pDC-SpCas9-hDHFR. The corresponding SpCas9 plasmid was used as a control. Plasmids were electroporated into synchronized early ring-stage parasites as previously described^42^. Following electroporation, transgenic parasites were maintained at 37°C and selected with 2.5nM WR99210 until stable transgenic parasite growth was re-established. The cultures were then divided into two groups and maintained at either 37°C or 28°C for 36 h. Genomic DNA was subsequently extracted, and HDR-mediated editing of the *gfp* locus was assessed by PCR using the primers listed in Supplementary Data 3.

---

**Extended Data Figure 1.**
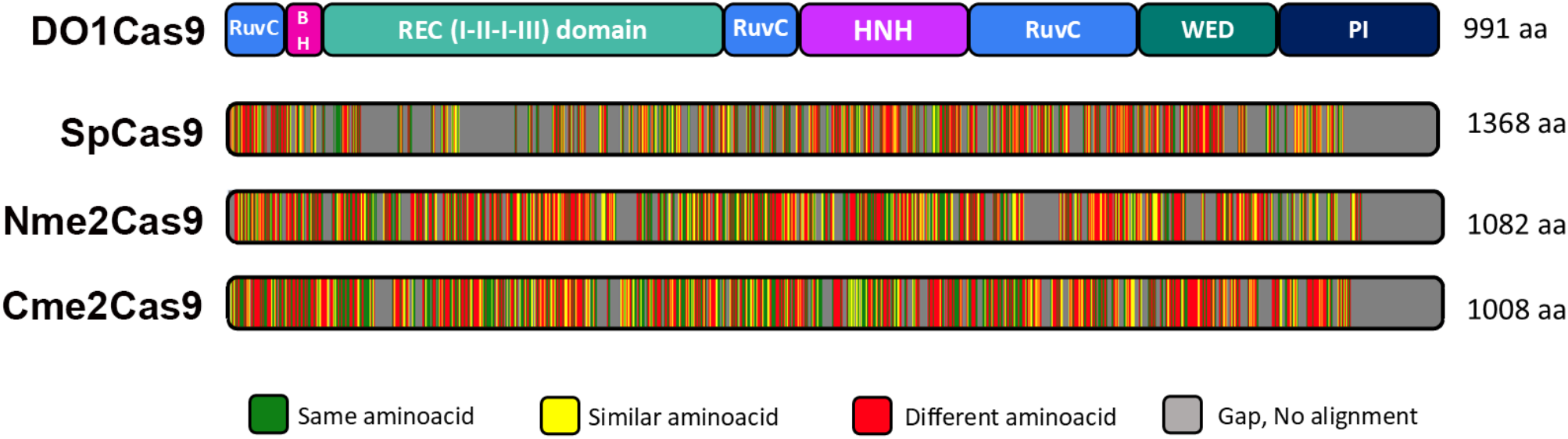
Alignment of protein sequences for DO1Cas9, SpCas9, Nme2Cas9 and Cme2Cas9 In silico pairwise protein sequence alignments between DO1Cas9, SpCas9 (Streptococcus pyogenes) (UniprotKB, Q99ZW2), Nme2Cas9 (Neisseria meningitidis) (Uniparc, UPI0001B02B97), and Cme2Cas9 (metagenomic Cas9 ortholog)^2^ sequences were performed using the EMBOSS needle algorithm1 (Needleman-Wunsch global alignment). Alignment outputs were subsequently processed and visualized using custom Python scripts. The domain architecture of DO1Cas9 is shown above the alignments, including the RuvC, bridge helix (BH), REC, HNH, WED, and PAM-interacting (PI) domains. Amino acid conservation is represented by colour coding: identical residues are shown in green, non-identical residues with similar biochemical properties in yellow, non-identical residues with dissimilar biochemical properties in red, and alignment gaps or unaligned positions in grey.

**Extended Data Figure 2.**
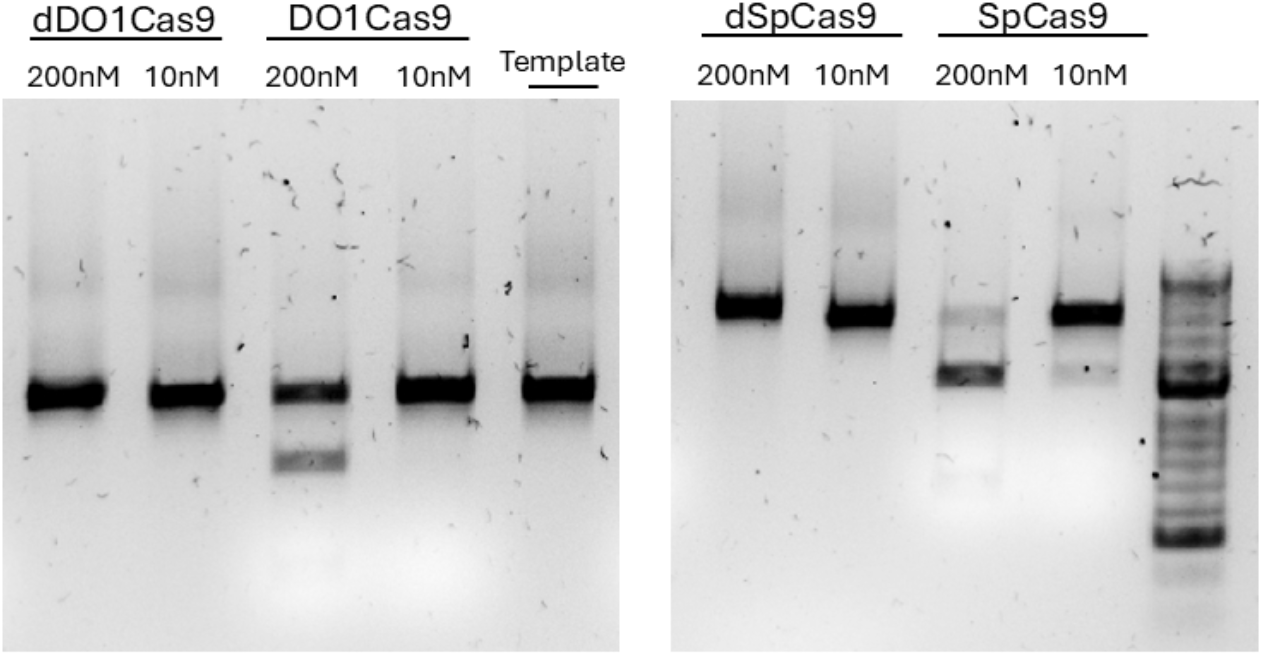
Validation of DO1Cas9 dead variant. In vitro cleavage activity of wild-type and nuclease-dead Cas9 variants. Ribonucleoprotein complexes containing wild-type DO1Cas9, dDO1Cas9 (D9A and H574A), SpCas9, or dSpCas9 (with mutations D10A and H840A, by IDT) were incubated with a target DNA substrate and analyzed by agarose gel electrophoresis. Cas9 concentrations of 200 nM and 10 nM were tested as indicated. Wild-type DO1Cas9 and SpCas9 generated the expected cleavage products, whereas no detectable cleavage was observed for dDO1Cas9 or dSpCas9, confirming that H574 and D9 are required for DO1Cas9 nuclease activity. Template, undigested 1456-bp PCR amplicon of the *Caenorhabditis elegans dpy-10* gene.

**Extended Data Figure 3.**
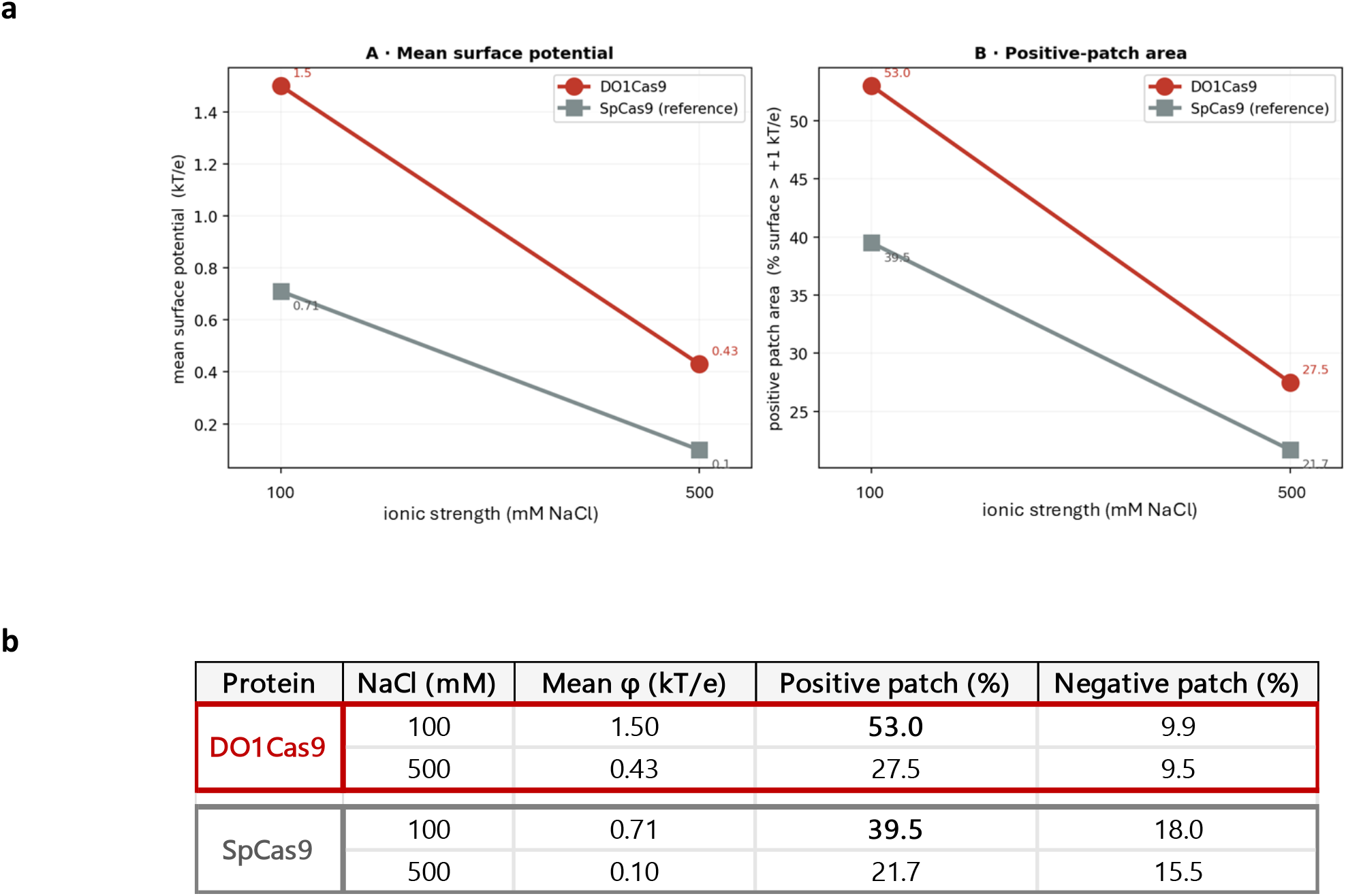
Surface electrostatic properties of DO1Cas9 and SpCas9. APBS (Adaptive Poisson-Boltzmann Solver) calculations performed on apo-state DO1Cas9 and SpCas9 structures at 100mM and 500mM NaCl. **a**, Mean surface electrostatic potential (ϕ, kT/e). **b**, Underlying electrostatic metrics obtained from the APBS analysis. Positive-patch area (surface fraction with ϕ > +1 kT/e). DO1Cas9 displays a markedly more electropositive surface than SpCas9 under both conditions. Increasing ionic strength attenuates surface electrostatic potential and positive-patch area, consistent with electrostatic screening.

**Extended Data Figure 4.**
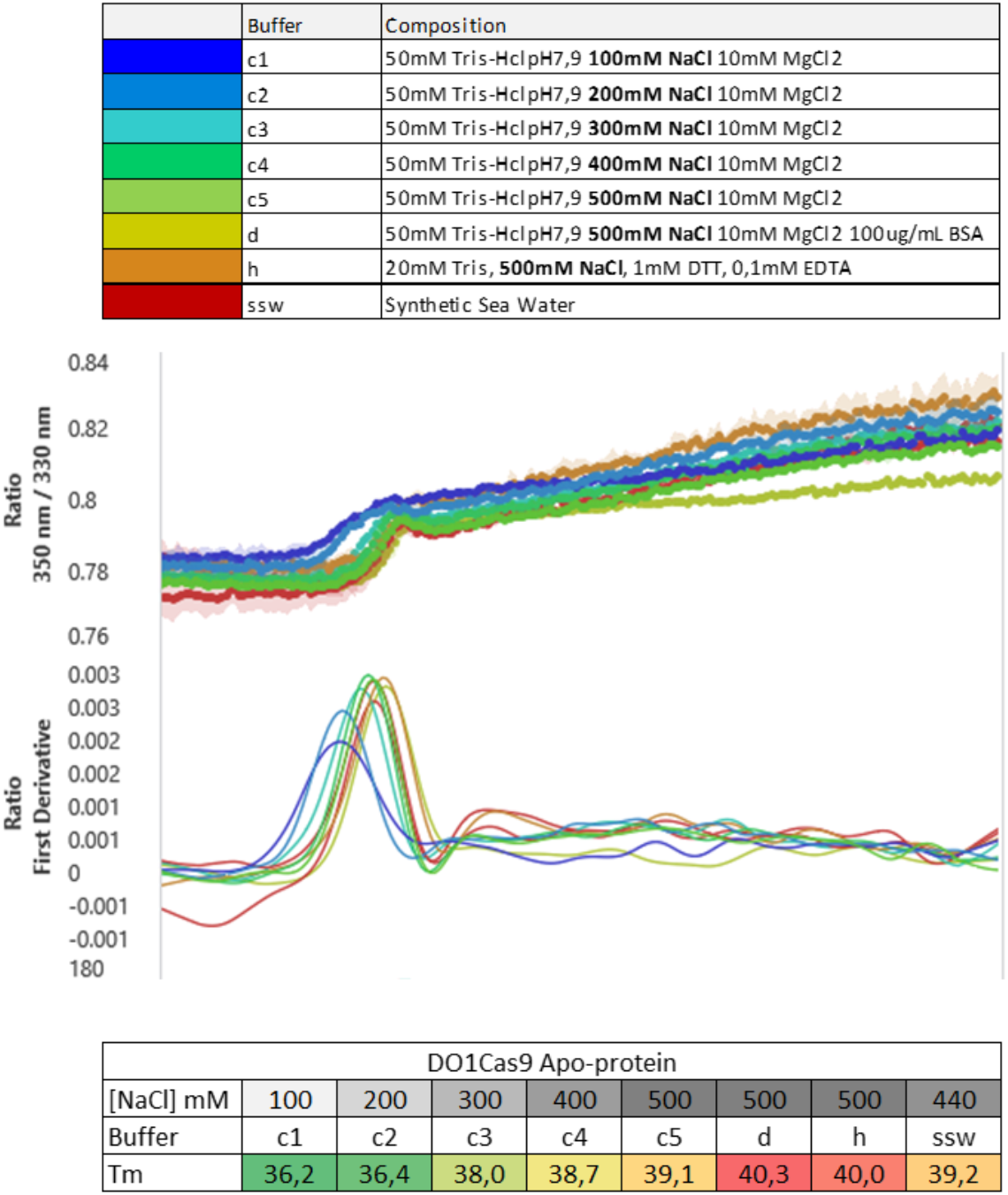
Influence of salt concentration in the thermal stability of apo DO1Cas9. Thermal stability of apo DO1Cas9 was assessed by nano-differential scanning fluorimetry (nanoDSF) across buffers containing increasing NaCl concentrations and seawater-mimicking conditions. Top panel, composition of buffers. Middle panel, intrinsic fluorescence ratio (F350/F330). Bottom panel, thermal transition midpoint (Tm) determined by first derivative Increasing ionic strength progressively shifted the thermal transition of apo DO1Cas9 towards higher temperatures, resulting in an overall increase in Tm of approximately 4°C between low-salt and high-salt conditions. Similar stabilization was observed in seawater-like buffers, consistent with the marine origin of DO1Cas9. These results indicate that elevated salt concentrations strongly stabilize the apo enzyme and support a model in which ionic-strength-dependent charge screening mitigates the destabilizing effects of the highly electropositive apo-state surface.

**Extended Data Figure 5.**
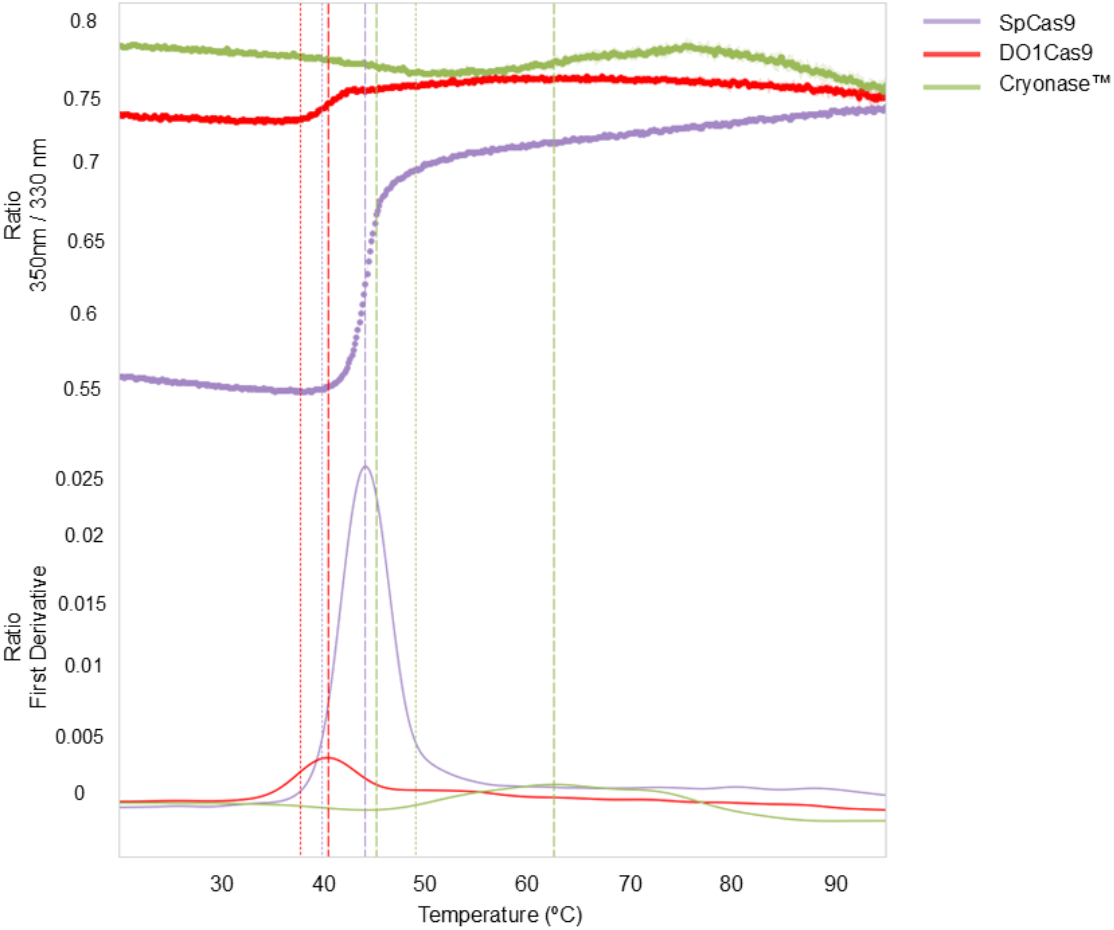
DO1Cas9, SpCas9 and Cryonase thermal-response profiles at nanoDSF. Thermal stability of the apo forms of DO1Cas9 (red), SpCas9 (purple), and the psychrophilic nuclease Cryonase™ (green) was analyzed by nanoDSF. The upper panel shows the intrinsic fluorescence ratio (F350/F330), and the lower panel shows the corresponding first derivative. Vertical dashed lines indicate the estimated unfolding onset temperature (T_on, upper panel) and thermal transition midpoint (T_m, lower panel). DO1Cas9 displayed a broad and gradual thermal-response profile closely resembling that of Cryonase™, whereas SpCas9 exhibited a sharp and cooperative thermal transition. These data indicate that DO1Cas9 shares biophysical characteristics commonly associated with cold-adapted enzymes while retaining the overall architecture of a Cas9 nuclease.

**Extended Data Figure 6.**
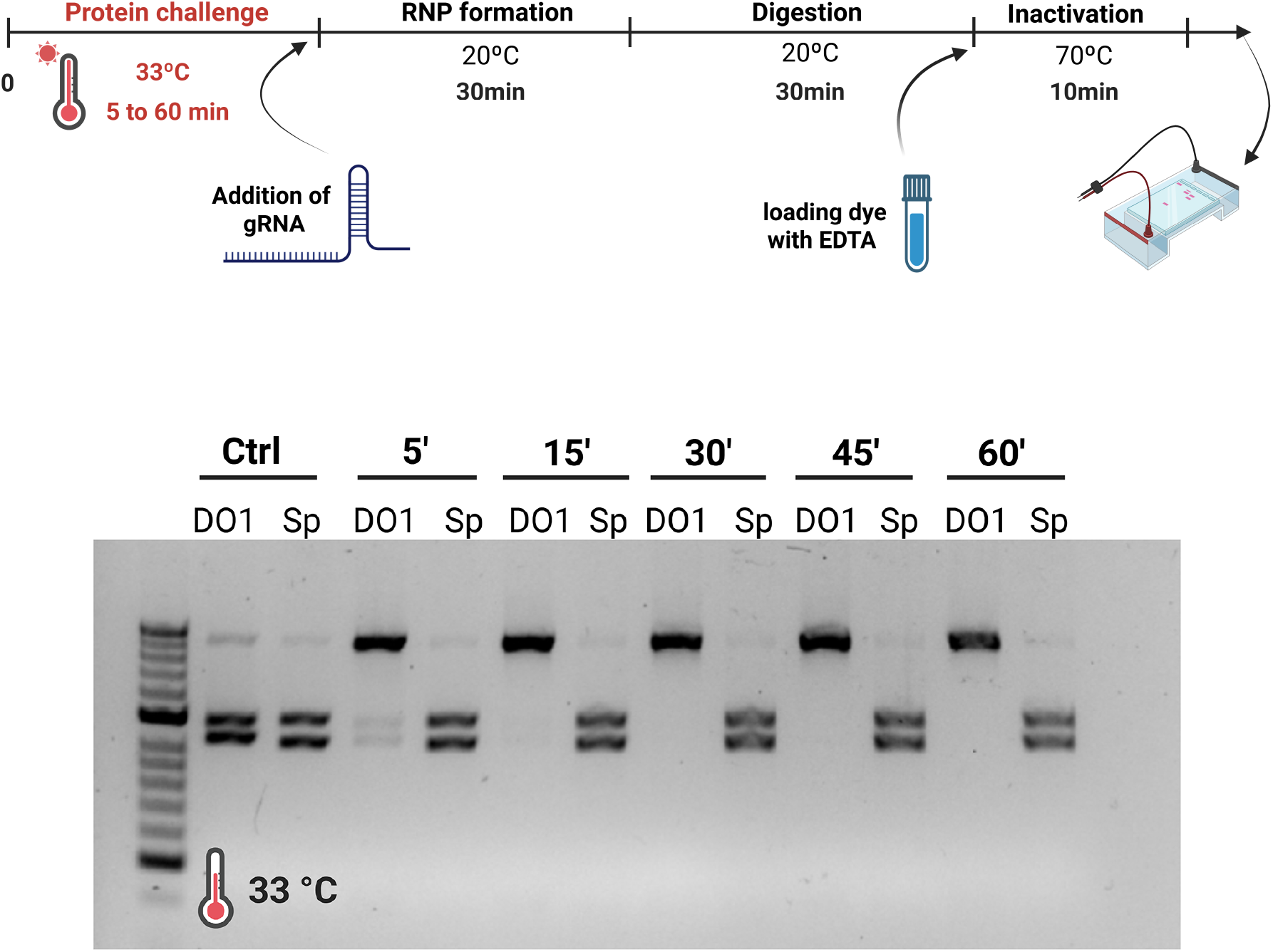
Thermal instability of DO1Cas9 apo-form at 33°C. Purified DO1Cas9 (DO1) or SpCas9 (Sp) proteins were pre-incubated at 33°C for the indicated times (5, 15, 30, 45, or 60 min) prior to guide RNA loading and ribonucleoprotein (RNP) assembly. Following RNP assembly, in vitro DNA cleavage assays were performed, and reaction products were analyzed by agarose gel electrophoresis. Control reactions (Ctrl) were assembled without prior heat treatment. A 15-min pre-incubation at 33°C markedly reduced DO1Cas9 cleavage activity, whereas SpCas9 remained active throughout the time course.

**Extended Data Figure 7.**
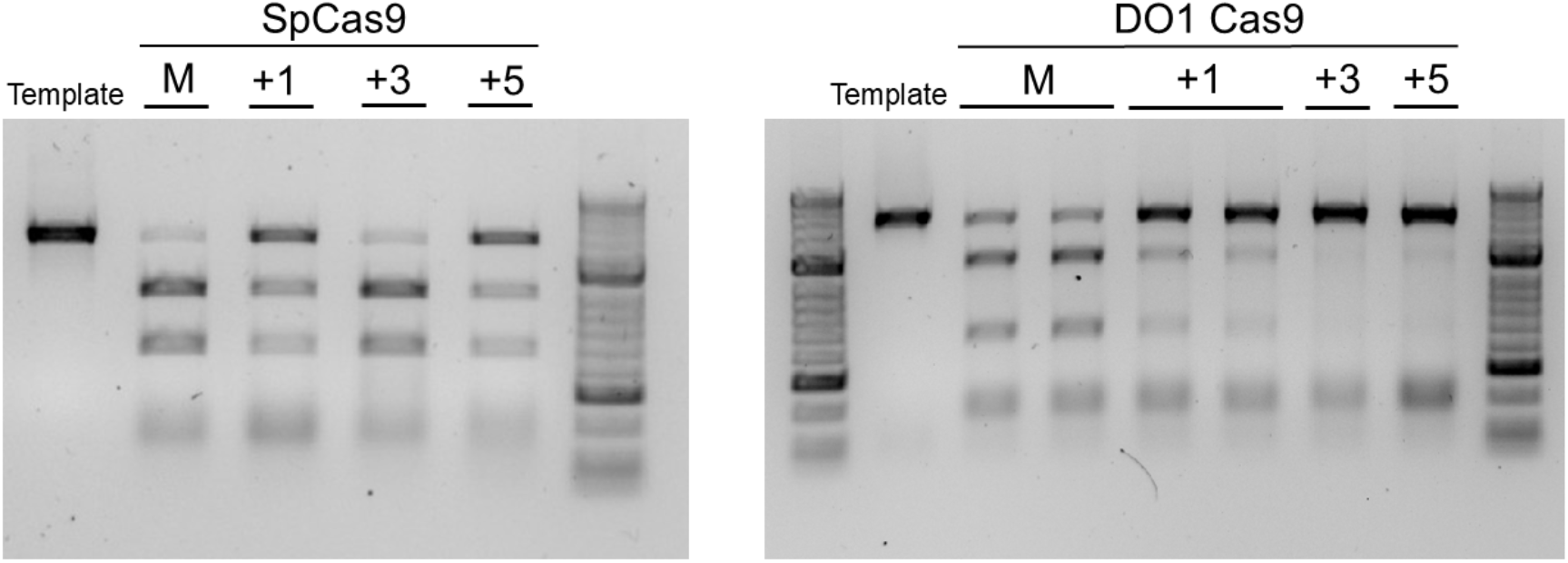
DO1Cas9 sensitivity to gRNA-target mismatches (*dpy-10* target). In vitro digestion of *dpy*-10 PCR amplicons by DO1Cas9 and SpCas9 using guide RNAs with either a perfectly matched spacer (M) or a single mismatch at positions +1, +3, or +5. Whereas SpCas9 cleaved substrates containing mismatches at all tested positions, DO1Cas9 showed markedly reduced cleavage activity in the presence of mismatches. Template, undigested *dpy-10* PCR amplicon control. Primers and gRNAs at Supplementary Data 3.

**Extended Data Figure 8.**
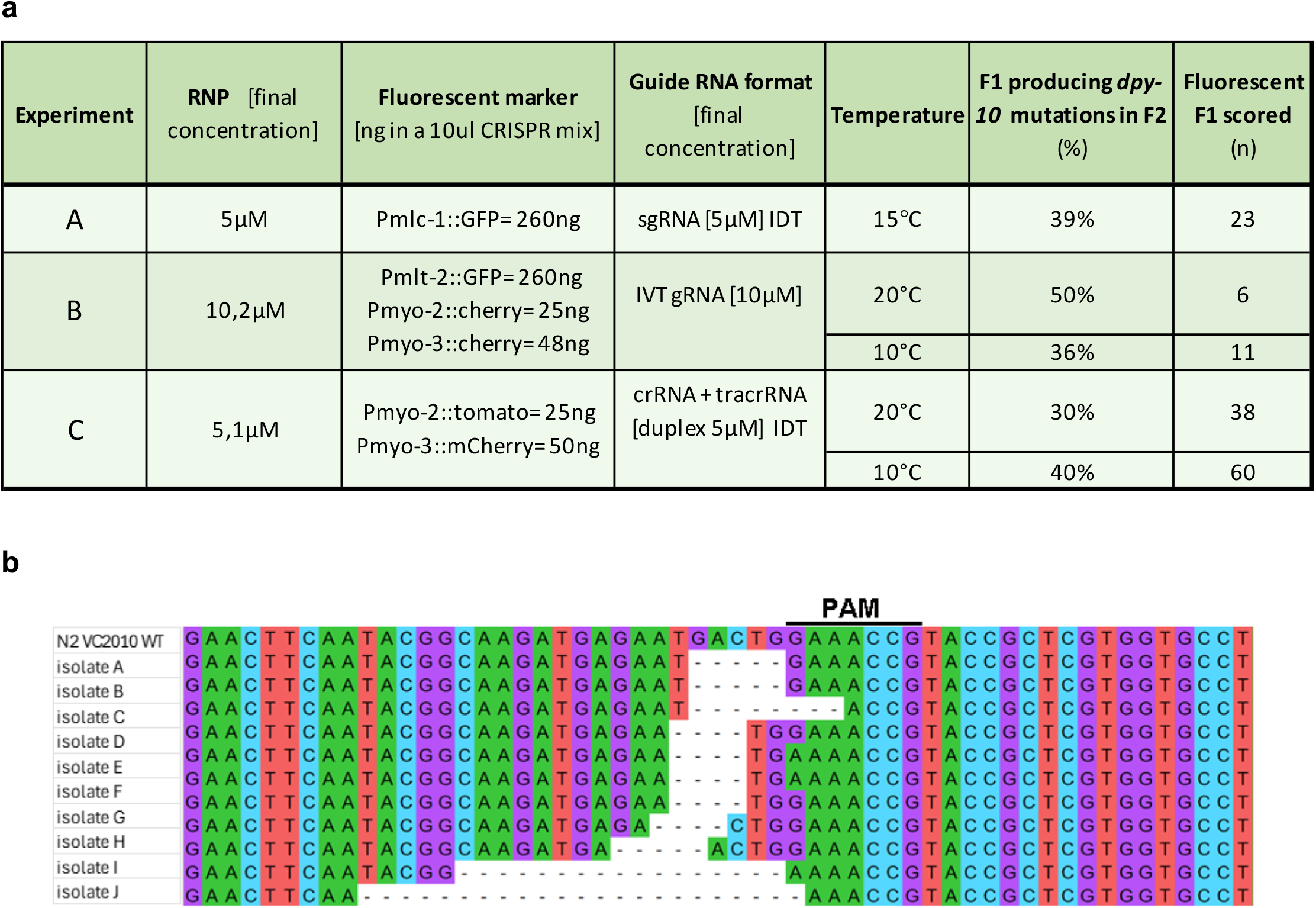
DO1Cas9 genome editing in *C. elegans*. **a**, CRISPR mixes summary of DO1Cas9 used in genome-editing experiments in *C. elegans*. Editing efficiency was calculated as the proportion of fluorescent F1 animals that produced *dpy-10* mutations in the F2 progeny, indicative of DO1Cas9 activity at the *dpy-10* locus. Temperature indicates temperature of worm culture after microinjection. The number of fluorescent F1 animals scored is indicated for each condition. **b**, Representative sequence alignment of mutations at the DO1Cas9-targeted *dpy-10* locus. Sanger sequences from independent edited individuals (isolates) homozygous for a *dpy-10* mutation (A-J) are aligned to the wild-type N2 VC2010 reference sequence. Dashes indicate deleted nucleotides. The NGG PAM is indicated above the corresponding sequence. Multiple insertion/deletion (indel) alleles were identified, demonstrating efficient generation of diverse repair outcomes following DO1Cas9-induced DNA cleavage in *C. elegans*.

**Extended Data Figure 9.**
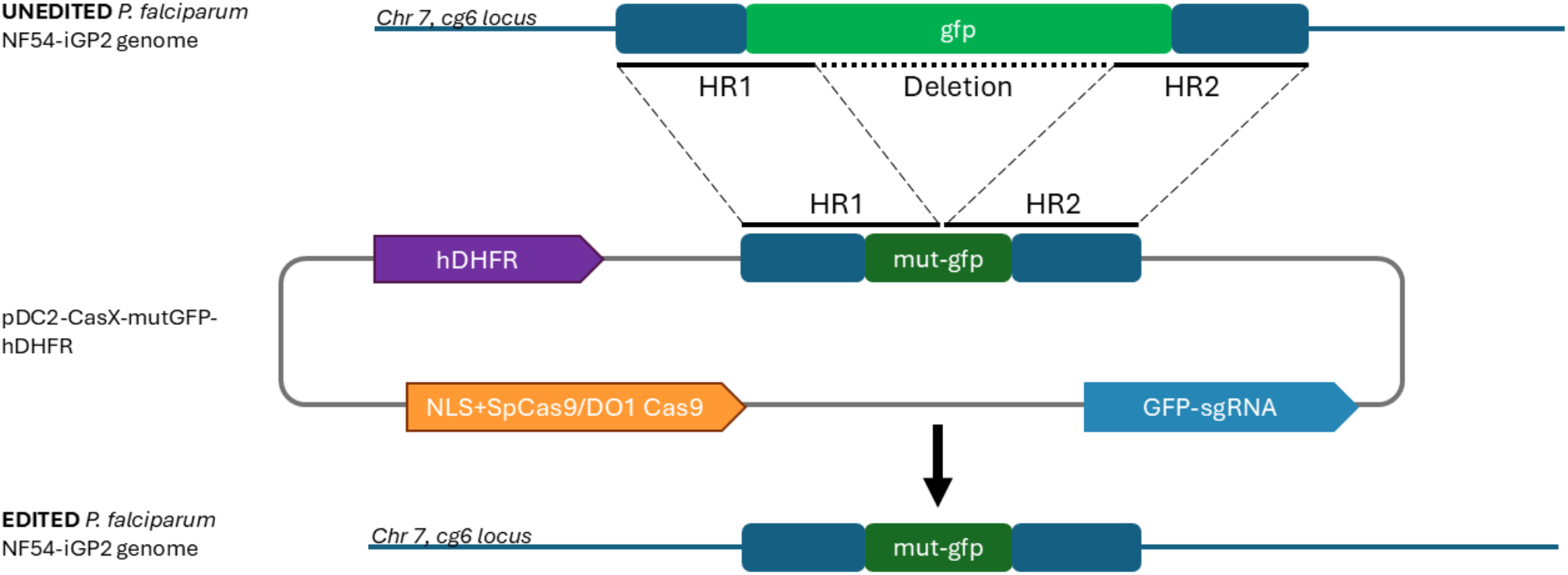
Schematic of the strategy for HDR-mediated genome editing of the gfp locus in Plasmodium falciparum using DO1Cas9 or SpCas9. Schematic of the homology directed repair (HDR)-based editing strategy used to evaluate DO1Cas9 activity in *P. falciparum* NF54-iGP2 parasites. The endogenous *gfp* reporter integrated at the chromosome 7 *cg6* locus was targeted with a gRNA directed against *gfp*. The editing plasmid encoded either DO1Cas9 or SpCas9 together with the *gfp*-targeting gRNA and a donor DNA template carrying a truncated *gfp* sequence (*mut-gfp*) flanked by homology regions (HR1 and HR2). Following Cas9-induced double-strand cleavage, repair by HDR replaced the endogenous gfp sequence with the shortened mut-gfp allele.

